# 2D or Not 2D? An FMRI Study of How Dogs Visually Process Objects

**DOI:** 10.1101/2020.06.04.134064

**Authors:** Ashley Prichard, Raveena Chhibber, Kate Athanassiades, Veronica Chiu, Mark Spivak, Gregory S. Berns

## Abstract

Given humans’ habitual use of screens, they rarely consider potential differences when viewing two dimensional (2D) stimuli and real-world versions of dimensional stimuli. Dogs also have access to many forms of screens and touch pads, with owners even subscribing to dog-directed content. Humans understand that 2D stimuli are representations of real-world objects, but do dogs? In canine cognition studies, 2D stimuli are almost always used to study what is normally 3D, like faces, and may assume that both 2D and 3D stimuli are represented in the brain the same way. Here, we used awake fMRI of 15 dogs to examine the neural mechanisms underlying dogs’ perception of two- and three-dimensional objects after the dogs were trained on either a two- or three-dimensional version of the objects. Activation within reward processing regions and parietal cortex of the dog brain to 2D and 3D versions of objects was determined by their training experience, as dogs trained on one dimensionality showed greater activation to the dimension on which they were trained. These results show that dogs do not automatically generalize between two- and three-dimensional stimuli and caution against implicit assumptions when using pictures or videos with dogs.

## INTRODUCTION

Studies of canine cognition frequently rely on two-dimensional (2D) pictures to test dogs’ ability to discriminate between objects, species, or faces (Albuquerque et al. 2016; Autier-Derian et al. 2013; Barber et al. 2016; Huber et al. 2013; Muller et al. 2015; Pitteri et al. 2014; Wallis et al. 2017). Visual stimuli for these studies are utilized because they are easy to obtain from studies on humans and nonhuman primates and are easy to implement in laboratory settings. But the ecological validity of this line of research hinges on the extent to which the findings transfer to real-world stimuli and contexts (Romero and Snow 2019). As dogs may not perceive 2D visual stimuli as humans do, are images appropriate stimuli for the study of dog cognition?

Visual stimuli are often selected without considering the nature of dogs’ visual perception (Miller and Murphy 1995). For example, dogs have a higher flicker fusion rate than humans. This means that they may perceive the flickering of a video display if the refresh rate is too low. With movies, dogs may notice a gap or flicker between frames. Dogs also have a visual streak as opposed to a fovea (as in primates), causing increased sensitivity to stimuli in the periphery of the visual field. This means that displaying a picture or playing a video to a dog may not accurately reflect what a dog sees in the real world because they may focus on different aspects of the video than we do (Byosiere et al. 2018; Byosiere et al. 2019). Although there is ample evidence that dogs can perceptually discriminate features of images, this does not mean that the images necessarily represent their real-world counterparts to the dog.

Research in canines and other nonhumans that utilize pictures share an underlying assumption that, like humans, dogs perceive 2D stimuli such as faces as similar to real 3D faces. Dogs do behaviorally differentiate pictures, as they can discriminate between pictures of human facial expressions and between pictures of familiar and strange dogs or humans (Autier-Derian et al. 2013; Barber et al. 2016; Huber et al. 2013; Muller et al. 2015; Pitteri et al. 2014; Somppi et al. 2012), and following substantial training, dogs show the ability to follow commands presented by humans through video projection (Pongracz et al. 2003). One study reported dogs’ use of duplicates of the objects or miniature versions as referents to retrieve the corresponding objects, concluding that dogs can use iconic representations (Kaminski et al. 2009). However, the same dogs did not perform well using pictures versions of the corresponding objects. Despite this widespread use of 2D visual stimuli in canine cognition, studies have not shown that dogs use 2D stimuli as referents for real world stimuli.

The ability to abstract from 2D to 3D versions of objects is not uniquely human. Many nonhuman species show evidence of behavioral transfer from pictures or videos to objects, pictures of food, or conspecifics following substantial training (Bovet and Vauclair 2000; Johnson-Ulrich et al. 2016; Wilkinson et al. 2013). This means that there is little evidence for 2D to 3D transfer happening naturalistically in a nonhuman species. Nor does recognition between pictures mean that the animal has abstract knowledge of objects, that they have formed a mental representation, or that they equate pictures and real world objects (Jitsumori 2010; Weisman and Spetch 2010).

Using functional magnetic resonance imaging (fMRI), regions of the primate brain have been identified as selective for processing specific types of visual stimuli, including the fusiform face area (FFA) for processing faces or the lateral occipital complex (LOC) for processing objects (Beauchamp et al. 2004; Durand et al. 2007; Eger et al. 2008; Janssen et al. 2018; Kourtzi and Kanwisher 2000; Kriegeskorte et al. 2008). Yet these fMRI studies have a similar caveat: they too rely on 2D visual stimuli as proxies for real-world stimuli and use subjects who are overly familiar with pictures.

There is some evidence that object processing regions of the human brain respond differently to 2D and 3D versions of stimuli. An fMRI study that directly compared neural activation within the LOC to real world objects and 2D versions of the same objects found that the LOC does not respond to the two versions of stimuli in the same way (Snow et al. 2011). In behavioral studies, real objects also prompt greater attention and memory retrieval than 2D images, and elicit goal-directed actions whereas 2D images do so to a lesser degree (Gomez et al. 2018; Snow et al. 2014). Goal directed actions, such as grasping, are difficult to generalize to 2D versions of objects because 2D versions lack the same binocular cues or proprioceptive feedback (Freud et al. 2018; Gallivan and Culham 2015; Hutchison and Gallivan 2018).

As in human studies of vision, fMRI can be used to elucidate the neural mechanisms underlying dogs’ perception of objects. FMRI studies of awake dogs have increased in complexity and duration, paralleling human fMRI studies. Canine studies show that stimulus-reward associations acquired prior to or during scanning are learned at different rates due to neural biases within the reward-processing regions of the brain, such as the caudate and amygdala (Cook et al. 2016; Prichard et al. 2018a). Dogs also process familiar human words associated with objects in similar language-processing regions of humans, like the temporal-parietal junction, and show greater activation to novel words versus familiar words (Prichard et al. 2018b). As in human imaging, functional localizers have also revealed areas of dogs’ occipital cortex selective for processing human and dog faces (Cuaya et al. 2016; Dilks et al. 2015; Szabo et al. 2020; Thompkins et al. 2018). Together these studies show that activation within areas of the dog brain can be used to predict perceptual or behavioral biases when processing visual stimuli.

In two separate studies, we used fMRI to measure activity in dogs’ brains in response to both objects and pictures of the objects. In Study 1, 15 dogs were split into two groups; dogs in the first group were trained on two 3D objects, and dogs in the second group were trained on two pictures of the objects. One stimulus was associated with reward and the other with nothing. During the fMRI session, dogs from both groups were presented with both the picture stimuli and object stimuli. If hedonic mechanisms facilitate abstraction from 2D to 3D (and vice-versa), then dogs should show greater neural activity in the caudate for the trained reward stimulus than the no reward stimulus, regardless of whether they were trained on objects or pictures. In Study 2, we developed a functional localizer for object processing regions analogous to LOC. If dogs equate 2D and 3D stimuli, then they should show no difference in neural activity between the object and the picture in these regions.

## MATERIALS AND METHODS

### Participants

Participants for both studies were 15 pet dogs volunteered by their Atlanta owners for fMRI training and fMRI studies (Prichard et al. 2018a) (Table 1). Each dog had previously completed two or more scans for the project and had demonstrated the ability to participate in MRI scans.

**Table 1.** Dogs (N=19) and participation in experiments.

| Dog | Breed | Sex | Object Localizer | 2D to 3D | S+ |
| --- | --- | --- | --- | --- | --- |
| Caylin | Border collie | F | 1 | 1 | 3D Giraffe |
| Daisy | Pitbull mix | F | 1 | 1 | 3D Giraffe |
| Eddie | Labrador Golden mix | M | 1 | 1 | 2D Whale |
| Kady | Labrador | F |  | 1 | 3D Whale |
| Koda | Pitbull mix | F | 1 | 1 | 2D Giraffe |
| Libby | Pitbull mix | F | 1 | 1 | 3D Whale |
| Mauja | Cattle dog mix | F |  | 1 | 2D Whale |
| Ohana | Golden Retriever | F | 1 | 1 | 3D Whale |
| Oliver | Border collie Beagle mix | M | 1 | 1 | 2D Whale |
| Pearl | Golden Retriever | F | 1 | 1 | 2D Giraffe |
| Tallulah | Carolina dog | F | 1 | 1 | 2D Whale |
| Truffles | Pointer mix | F | 1 | 1 | 3D Giraffe |
| Tug | Portuguese Water dog | M |  | 1 | 3D Giraffe |
| Zen | Labrador Golden mix | M | 1 | 1 | 2D Giraffe |
| Zoey | Goldendoodle | F | 1 | 1 | 3D Giraffe |
| **N** |  |  | **12** | **15** |  |

Dog's names, breed, sex, and participation in experimental studies (indicated by 1) are listed. Grey denotes a dog participated in all 3 studies.

### General Experimental Design

The experimental design was similar to previous dog fMRI studies that examined preference using visual stimuli associated with food or social reward (Cook et al. 2016). Briefly, dogs entered and positioned themselves in custom chin rests in the scanner bore. All scans took place in the presence of the dog’s owner, who stood out of view of the dogs throughout the scan near the opening of the scanner bore and delivered all rewards (hot dogs) to the dog. An experimenter was stationed next to the owner, out of view of the dog, where the experimenter controlled the presentation of stimuli to the dogs. The onset and offset of each stimulus were timestamped by the simultaneous press of a four-button MRI-compatible button box by the experimenter.

### Imaging

Scanning was conducted with a Siemens 3 T Trio whole-body scanner using procedures described previously (Berns et al. 2013; Berns et al. 2012; Prichard et al. 2018a; Prichard et al. 2018b). The functional scans used a single-shot echo-planar imaging (EPI) sequence to acquire volumes of 22 sequential 2.5 mm slices with a 20% gap (TE = 25 ms, TR = 1260 ms, flip angle = 70°, 64 × 64 matrix, 2.5 mm in-plane voxel size, FOV = 192 mm). Slices were oriented dorsally to the dog’s brain (coronal to the magnet, as in the sphinx position the dogs’ heads were positioned 90 degrees from the prone human orientation) with the phase-encoding direction right-to-left. Four runs of up to 400 functional volumes were acquired for each subject, with each run lasting about 9 minutes. Following functional scans, a T2-weighted structural image of the whole brain was acquired using a turbo spin-echo sequence (25-36 2mm slices, TR = 3940 ms, TE = 8.9 ms, flip angle = 131°, 26 echo trains, 128 x 128 matrix, FOV = 192 mm).

### Preprocessing

Preprocessing was the same as described in previous studies (Berns et al. 2013; Prichard et al. 2018a). Briefly, preprocessing of the fMRI data included motion correction, censoring, and normalization using AFNI (NIH) and its associated functions. A hand-selected reference volume for each dog that corresponded to their average position within the magnet bore across runs was used for two-pass, six-parameter rigid-body motion correction. Aggressive censoring removed unusable volumes from the fMRI time sequence because dogs can move between trials and when consuming rewards. Data were censored when estimated motion was greater than 1 mm displacement scan-to-scan and also based on outlier voxel signal intensities. A mask was drawn in functional space for each dog in the cerebellum, which was used to censor the data further by removing volumes where the beta values extracted from the cerebellum were assumed to be beyond the physiologic range of the BOLD signal (> |3 percent signal change|) for each trial. Smoothing, normalization, and motion correction parameters were identical to those described in previous studies (Prichard et al. 2018a). EPI images were smoothed and normalized to %-signal change with 3dmerge using a 6mm kernel at full-width half-maximum. The Advanced Normalization Tools (ANTs) software was used to spatially normalize the mean of the motion-corrected functional images (Avants et al. 2011) to the individual dog’s structural image. We also performed a nonlinear transformation from each dog’s structural image to a high-resolution canine brain atlas, developed from a previous study of Labrador retrievers (Berns et al. 2017).

### Experimental Design

#### Study 1: 2D vs. 3D

In each session, dogs were presented with two objects (a stuffed giraffe and a stuffed whale) and two life-sized cut-out pictures posted on foamboard of the objects (Fig 1). Each stimulus was attached to a three-foot dowel that the experimenter used to present the stimuli to the dog while inside the scanner bore. Neither object had been encountered before by the dogs. Dogs were semi-randomly split into two groups prior to scanning, 8 in the object group and 7 in the picture group. Prior to the first run, dogs were trained on the stimulus-reward associations (10 reward, 10 no-reward) based on their assigned group. Dogs were also refreshed on the stimulus-reward associations between runs (5 reward, 5 no-reward). Following each run, dogs would exit the scanner and rest or drink water.

**Figure 1.**
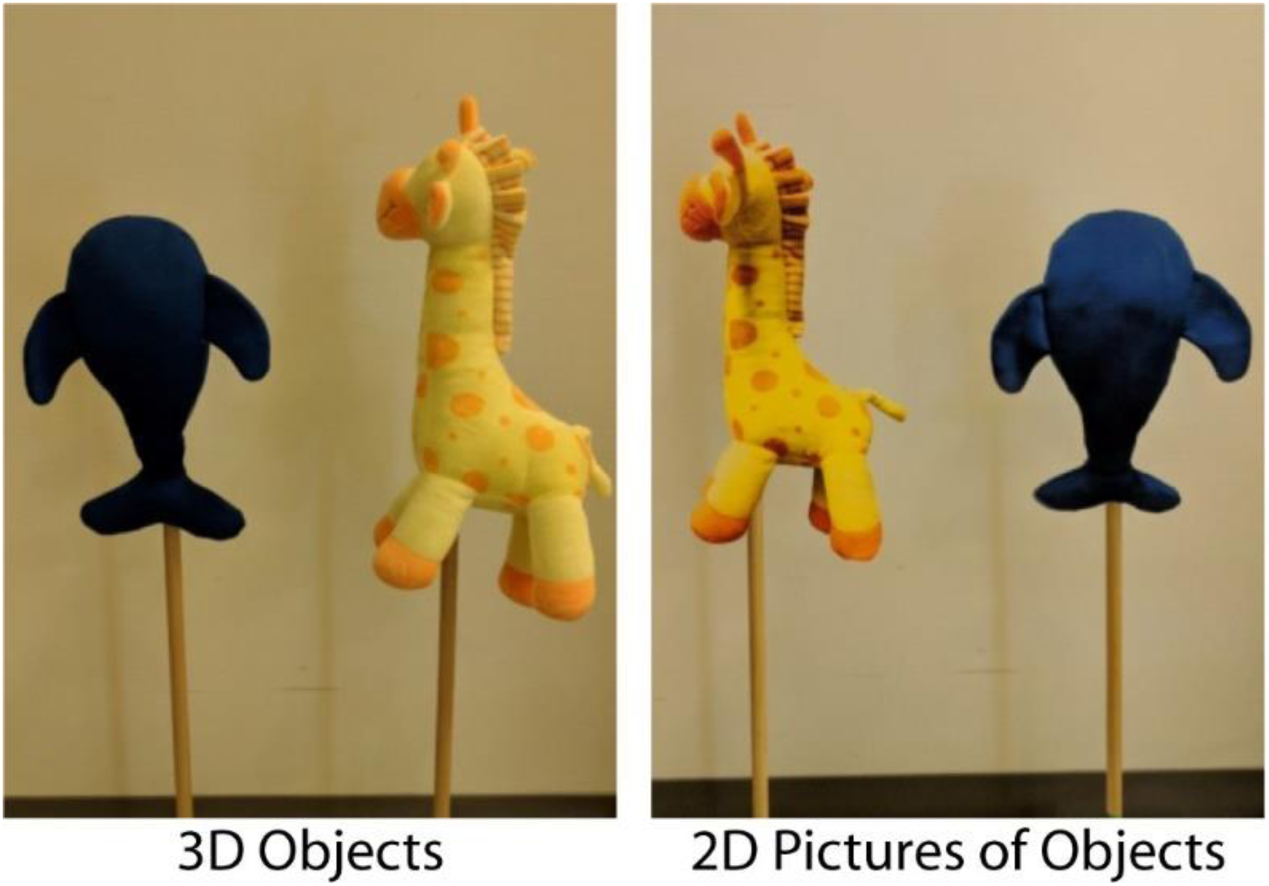
2D & 3D Stimuli. **Left)** 3D whale and 3D giraffe objects attached to 2.5-foot dowels for presentation of stimuli to dogs while in the scanner. **Right)** Pictures of the whale and giraffe 3D objects were printed to create 2D color-matched versions of the 3D stimuli and pasted to foam board and 2.5-foot dowels for presentation of 2D stimuli to dogs while in the scanner

Dogs in the object group were trained on object stimuli and were semi-randomly assigned the whale or giraffe as the reward stimulus. The presentation of the reward object (giraffe or whale) was immediately followed by the delivery of a food reward, and presentation of the other object was immediately followed by nothing. In the picture group, dogs were trained on picture stimuli and were semi-randomly assigned the whale or giraffe as the reward stimulus. The presentation of the reward picture (giraffe or whale) was immediately followed by the delivery of a food reward, and the other picture was immediately followed by nothing. Training on the conditioned stimuli occurred prior to each run when the dog was positioned in the scanner bore, but before scan acquisition. During scan acquisition, no stimuli were followed by the delivery of a food reward, so that dogs could not discriminate between objects and pictures based solely on food reward. To maintain general motivation to stay in the scanner, food rewards were presented by the owner randomly throughout the scan session between presentations of the stimuli.

An event-based design was used, consisting of trained reward and trained no-reward trial types, as well as symbolic reward and symbolic no-reward trial types. Trained reward and trained no-reward trials consisted of the two conditioned stimuli associated with food reward prior to scanning (e.g. objects for half of the dogs, pictures for the other half). Symbolic reward and symbolic no-reward trials consisted of the two untrained stimuli (e.g. pictures for dogs trained on objects, and objects for dogs trained on pictures). On all trials, a stimulus was presented for a 5 s duration, followed by nothing. Trials were jittered to randomize presentation order and were separated by a variable inter-trial interval. Each dog received the same trial sequence.

A scan session consisted of 4 runs, lasting approximately 9 minutes per run. Each run consisted of 25 trials (5 trained reward, 5 trained no-reward, 5 symbolic reward, 5 symbolic no-reward, and 5 food rewards delivered at random), for a total of 100 trials per scan session. No trial type was repeated more than 4 times sequentially, as dogs could habituate to the continued presentation of a stimulus.

### Analyses

Each subject’s motion-corrected, censored, smoothed images were analyzed within a general linear model (GLM) for each voxel in the brain using 3dDeconvolve (part of the AFNI suite). Task related regressors for each experiment were modeled using AFNI’s dmUBLOCK and stim_times_IM functions and were as follows: (1) trained reward stimulus; (2) trained no-reward stimulus; (3) symbolic reward stimulus; (4) symbolic no-reward stimulus. This function created a column in the design matrix for each trial, allowing for the estimation of beta values for each trial. Data were censored for outliers as described above for the contrasts of interest.

A series of contrasts were pre-planned to assess main effects related to the acquisition of trained associations and whether they generalized between 2D and 3D versions. Acquisition of the trained stimulus-reward association was probed with the contrast [trained reward— trained no reward]. Transfer of the trained reward and no-reward association to the untrained stimuli was probed with the contrast [symbolic reward—symbolic no reward]. A direct association between the trained and untrained reward stimuli was tested with the contrast [trained reward – symbolic reward]. The contrast [all_3D—all_2D] was performed to test for perceived differences between all 3D and all 2D stimuli, regardless of training. The average difference between trained stimuli and symbolic stimuli was assessed with the contrast [(trained reward + trained no-reward)—(symbolic reward + symbolic no-reward)]. Finally, the interaction between reward and no reward stimuli and symbolism was measured with the contrast [(trained reward — trained no reward)—(symbolic reward —symbolic no reward)].

### Region of Interest (ROI) Analysis

#### Caudate

As our interest was based on the dog’s response to trained stimuli versus symbolic stimuli, quantitative analyses based on the imaging results used activation values in the canine brain area previously observed to be responsive to reward stimuli (Cook et al. 2016). Anatomical ROIs of the left and right caudate nuclei were defined structurally using each dog’s T2-weighted structural image. ROI-based analyses were performed in individual, rather than group space.

Beta values for the contrasts comparing the change in activation to reward and no reward stimuli for trained (20 reward trials, 20 no-reward trials) and symbolic stimuli (20 reward trials, 20 no-reward trials) were extracted from the caudate ROIs in the left and right hemispheres. Beta values greater than an absolute four percent signal change were removed prior to analyses (assuming that these were beyond the physiologic range of the BOLD signal). The remaining beta values were analyzed using the mixed-model procedure in SPSS 24 (IBM) with fixed-effects for the intercept, group, hemisphere (left or right), and contrast type, identity covariance structure, and maximum-likelihood estimation.

#### Whole Brain Analysis

Each subject’s individual-level contrast from the GLM was normalized to the Labrador Retriever atlas space via the ANTs software. Spatial transformations included a rigid-body mean EPI to structural image, affine structural to template, and diffeomorphic structural to template. These spatial transformations were concatenated and applied to individual contrasts from the GLM to compute group level statistics. 3dttest++, part of the AFNI suite, was used to compute a t-test across dogs against the null hypothesis that each voxel had a mean value of zero. All contrasts from the GLM mentioned above were included. The average smoothness of the residuals from each dog’s time series regression model was calculated using AFNI’s non-Gaussian spatial autocorrelation function 3dFWHMx –acf. The acf option leads to greatly reduced FPRs clustered around 5 percent across all voxelwise thresholds (Cox et al. 2017). AFNI’s 3dClustsim was then used to estimate the significance of cluster sizes across the whole brain after correcting for familywise error (FWE). Similar to human fMRI studies, a voxel threshold of p ≤ 0.005 was used, and a cluster was considered significant if it exceeded the critical size estimated by 3dClustsim for a FWER ≤ 0.05, using two-sided thresholding and a nearest-neighbor of 1.

### Study 2: Object Localizer

To identify object-processing regions, dogs were presented with 3-s color movie clips projected on a screen in the bore of the magnet. Videos included human faces, novel objects (toys), familiar objects, and scram-bled objects (a 15 by 15 box grid with spatially rearranged movie frames.) Stimuli were presented using Python 2.7.9 and the Psychopy Experiment library. A blocked fMRI design was used where each block was 21 s with seven movie clips for each category. Each run contained two sets of four consecutive stimulus blocks in palindromic order. Stimulus blocks had a delay of 10 s, where dogs were fed intermittently between blocks, such that each run was approximately 7 minutes. On average, each dog completed three runs.

#### Analyses

As in Study 1, a general linear model was estimated for each voxel using AFNI’s 3dDeconvolve. Task related regressors were: (1) faces, (2) novel objects, (3) trained objects, and (4) scrambled objects. Individual object-specific regions, such as LOC, were identified with the contrast [novel objects—faces]. Each dog’s object-specific region was defined by the voxel threshold of the statistical map for the [novel objects—faces] contrast until the number of voxels in each ROI was approximately 40 voxels or less (Aulet et al. 2019). Beta values from the contrasts of interest mentioned in Study 1 were extracted from the object-specific region of each dog to examine potential differences in neural activation between 2D and 3D objects from the contrasts mentioned above.

## RESULTS

### Study 1: 2D vs 3D Results

#### Caudate ROI Analyses

There was differentiation of the reward and no-reward stimuli in the caudate ROIs for the trained stimuli, regardless of whether dogs were trained on objects or pictures. There was also a significant interaction of training x [Reward – No Reward] (*F* (1,45) = 11.29, *p* = 0.002) (Fig 2). This indicates that the trained reward association did not transfer to the symbolic stimuli.

**Figure 2.**
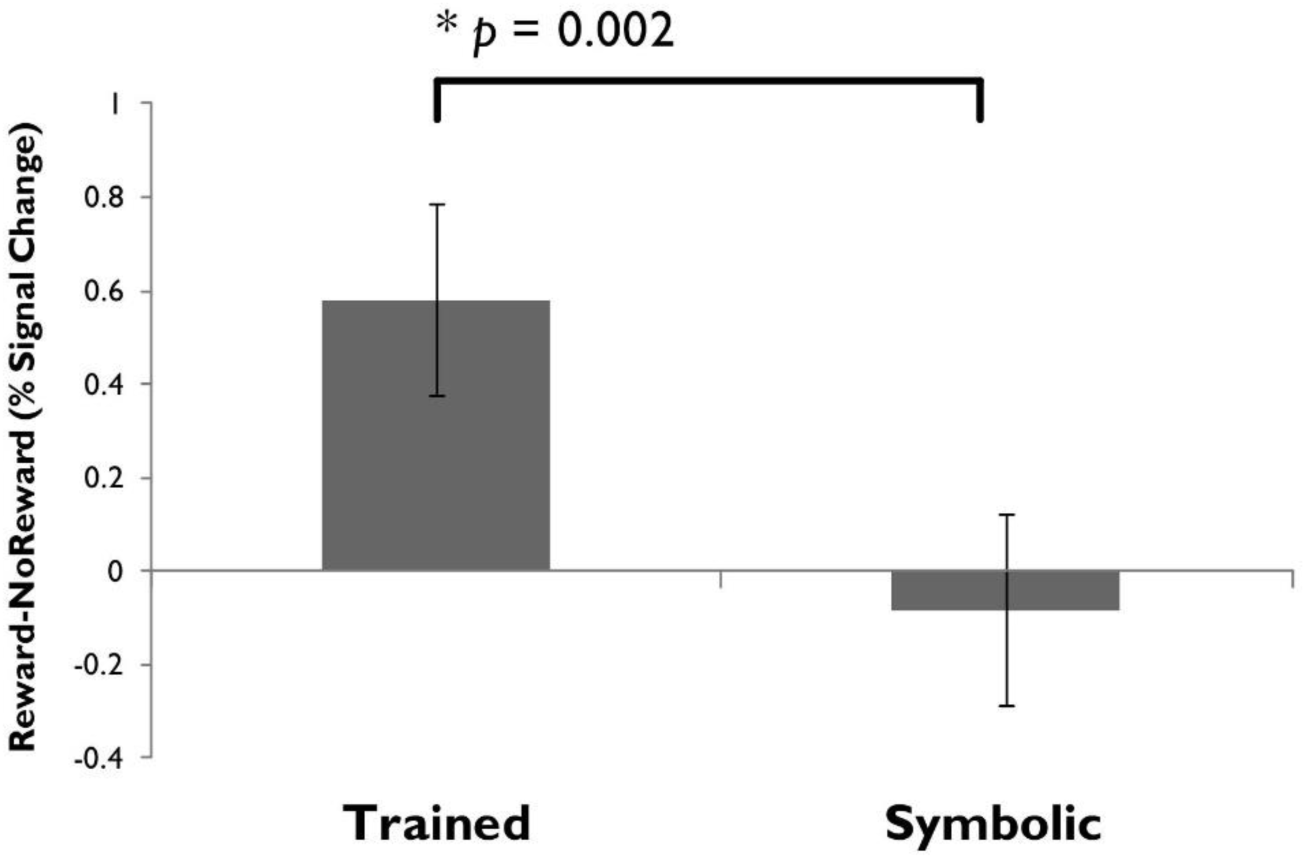
Average beta values (Percent signal change) in individual dogs’ caudate nucleus for the contrast of Reward— No Reward separated by training and testing (symbolic) stimuli. Changes in brain activation were extracted from contrasts in the 2D to 3D study. In the caudate there was a significant interaction of training x [Reward – No Reward] (*F* (1,45) = 11.29, *p* = 0.002)

#### Whole Brain Analyses

We found neural evidence for the differentiation of stimuli as an effect of the dimensionality of the training stimuli. Whole brain analysis of the contrasts of interest revealed significant activation for three contrasts (Table 2). The [trained reward—symbolic reward] contrast and the contrast comparing activation to the trained stimulus dimensionality versus the untrained stimulus dimensionality [trained reward + trained no-reward)—(symbolic reward + symbolic no-reward)] revealed a region in the posterior parietal lobe with greater activation toward the trained dimensionality of stimuli than the untrained dimensionality (Fig 3A). The contrast comparing the reward associations for the untrained dimension of stimuli [symbolic reward— symbolic no-reward] revealed a region in the right anterior parietotemporal cortex (Fig 3B).

**Table 2.** Cluster size and threshold significance for 2D and 3D object processing regions.

| 2D3D Contrast | Region | Voxel threshold | Cluster Size (1 mm<sup>3</sup> voxels) |
| --- | --- | --- | --- |
| trained reward – symbolic reward | Left posterior parietal | .005 | 454 (p<.02) |
| symbolic reward— symbolic no-reward | Right parietotemporal | .005 | 248 (p=.05) |
| (trained reward + trained no-reward)—<br>(symbolic reward + symbolic no-reward) | Left posterior parietal | .005 | 528 (p<0.01) |

**Figure 3.**
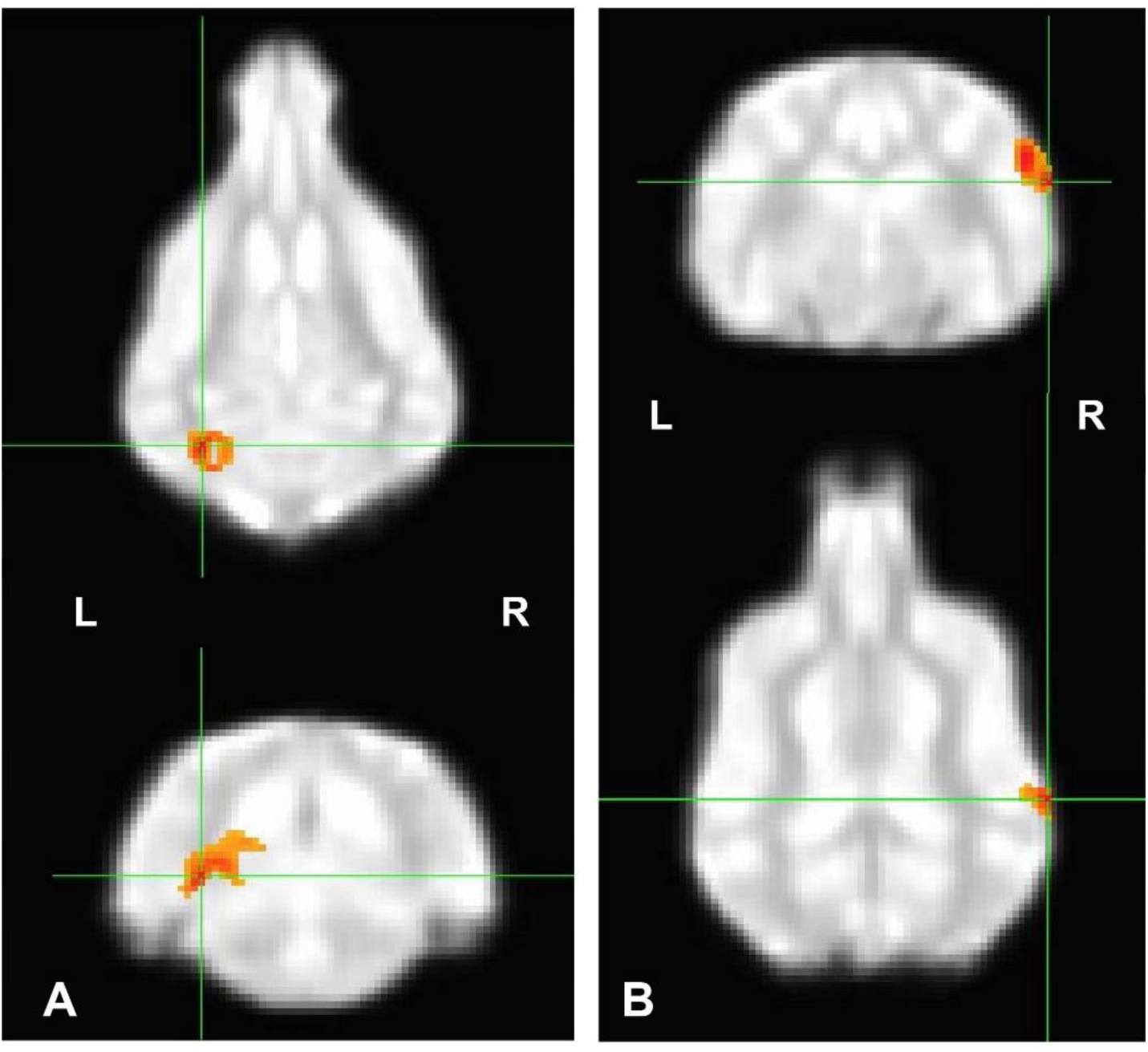
Regions important for the discrimination of dimensional object stimuli. Whole brain analysis of the contrasts of interest revealed significant activation only for three contrasts that survive a voxel threshold of 0.005. **A) The contrast comparing the trained S+ to the corresponding untrained dimension [trained reward— symbolic reward]** (454 voxels) revealed a region in the left posterior parietal lobe with greater activation toward the trained dimensionality of stimuli. **B) The contrast comparing the untrained dimension of stimuli [symbolic reward— no-reward]** (248 voxels) produced a region in the right anterior parietotemporal cortex.

### Study 2: Object Localizer Results

#### Individual Object Regions

Three dogs (Velcro, Rookie, and Zoey) failed to complete three runs of the object localizer task such that there was insufficient data to localize object-specific regions in the brain. Twelve dogs had object selective regions defined by the contrast [novel_objects—faces] in overlapping regions in either the left or right hemisphere (Fig. 4). We further examined these object regions for differences between 3D and 2D versions of the objects from the contrasts in Study 1. However, there were no statistically significant results for any of the contrasts in the object-specific regions across dogs.

**Figure 4.**
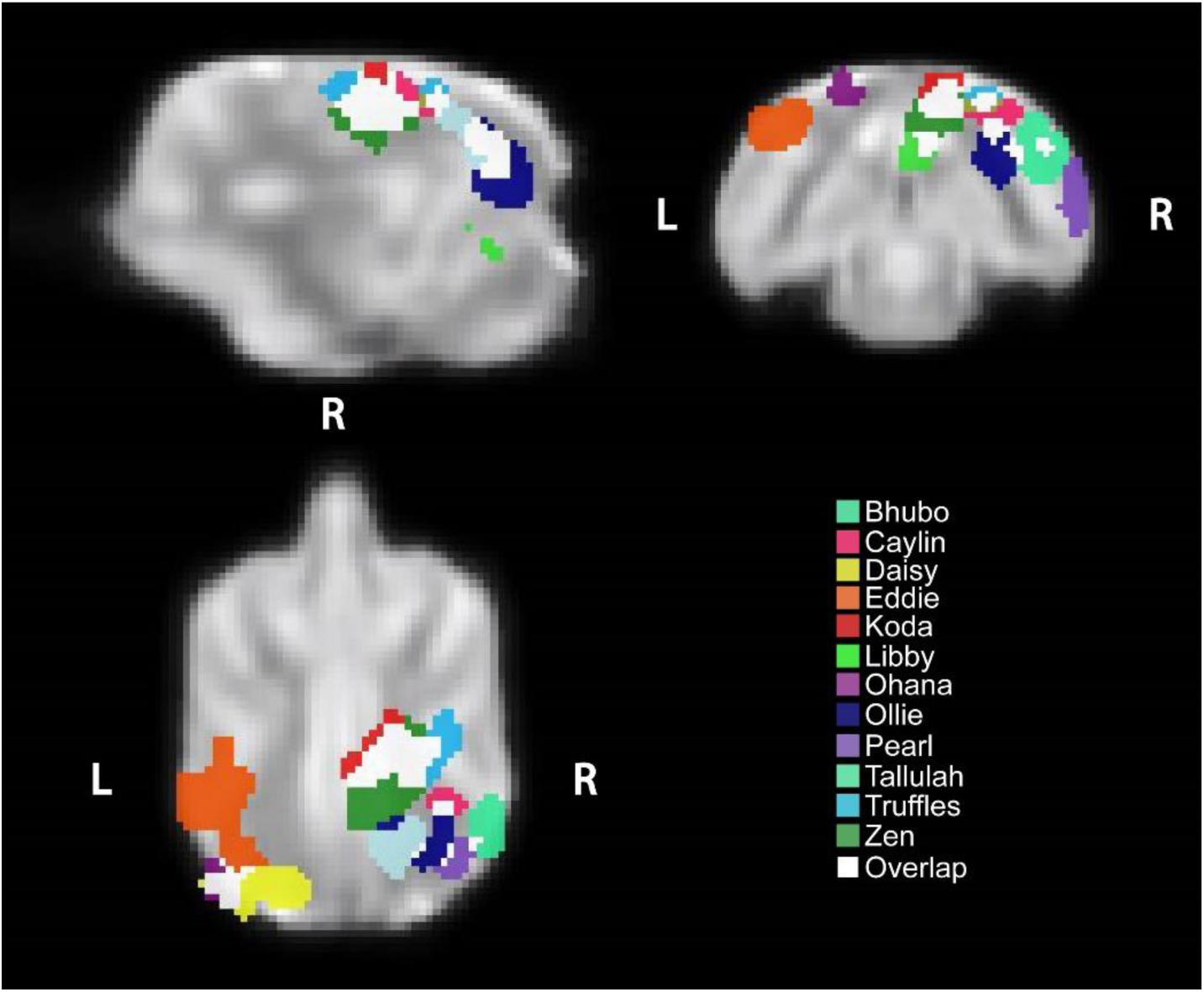
Individual Dog Object Regions. Sagittal, transverse, and dorsal sections. Regions were defined using the objects-faces contrast of video stimuli for each dog. Colors represent individual dogs; white represents overlap between two or more dogs.

## DISCUSSION

Our fMRI results provide the first evidence for neural differences in the occipital and parietal cortices of the dog brain for the processing of two- and three-dimensional objects. The main finding is that dogs’ perception of 2D and 3D objects is influenced by their experience with either stimulus dimension. Activation within reward processing regions was greater for the dimensionality of the trained reward stimulus. Whole-brain analyses revealed a left posterior parietal region selective for the trained dimension of stimuli over the untrained dimension. Taken together, these findings suggest that the neural representation of objects depends on dogs’ familiarity with two- and three-dimensional objects.

### Object Regions

In humans, viewing real objects as well as images of objects activates similar networks, particularly the lateral occipital complex along the lateral and ventral convexity of occipito-temporal cortex (Snow et al. 2011; Todd et al. 2012). However, in a human fMRI study that presented real objects and pictures of the objects, the LOC in particular was sensitive to visual differences between the two, such that LOC did not code the real (3D) and pictorial (2D) versions of a shape as equivalent (Snow et al. 2011). Because real objects afford specific actions, including the graspability of an object or if it is within reach of the dominant hand, object-specific actions may have a unique effect on neural responses to the different versions of the same object stimuli (Gallivan et al. 2009; Gallivan and Culham 2015; Gallivan et al. 2011; Snow et al. 2014). Unlike humans, we found little difference in dogs’ neural activation in individual object regions between 2D and 3D versions of object stimuli associated with reward. Our finding of similar neural activation in object regions of the dogs’ brains to 2D and 3D versions of object stimuli could be due to dogs forming an abstract object concept that is invariant to the dimensionality of the object. However, as the object-reward pairings were acquired using a passive viewing task, dogs had little experience interacting with the objects or picture versions of the objects. The dogs’ lack of action-associations with either object may therefore have made both objects and pictures of objects equivalent to the dog as neither was actionable. It is also possible that the study was insufficiently powered to detect potentially smaller effect sizes in dogs than in humans.

### Dimensionality Regions

As most studies of canine cognition rely on visual stimuli, we examined whether dogs use hedonic neural mechanisms to generalize from pictures of objects to real world objects. In the interaction contrast, we found that dogs show greater activation within the caudate nucleus to the trained dimension of stimuli relative to the untrained dimension (e.g. dogs trained on pictures of objects had greater activation to pictures relative to real world objects), suggesting hedonic neural mechanisms are biased toward the dimensionality of stimuli with which they are more familiar. Additional brain regions selective for stimulus dimensionality included a left posterior parietal region across dogs where there was greater activation to the trained dimensionality of stimuli than to the untrained dimensionality, which appeared in the same region but opposite hemisphere as the LOC defined in each dog in the object localizer study. Multi-voxel pattern analysis (MVPA) of human imaging data supports these findings, as patterns in object regions can be different for object exemplars from the same category that vary based on viewpoint or size, as well as between 2D and 3D versions of the same objects (Eger et al. 2008; Snow et al. 2011). Consistent with human imaging studies, the left posterior region also showed greater activation to objects relative to faces across dogs and appeared in regions of the canine brain similar to the primate LOC (Freud et al. 2017; Freud et al. 2018).

There was also greater activation to the untrained reward versus no reward stimuli in a right parietotemporal region across dogs (e.g. dogs trained that the 2D giraffe was the reward stimulus had greater activation to the 3D giraffe than the 3D whale in this region). Greater activation to the untrained reward stimulus in this region provides some evidence that dogs use hedonic neural mechanisms to generalize a stimulus-reward association from the trained reward stimulus to the untrained stimulus. However, we do not know what features, such as color or shape, that the dog may use to facilitate this representation. In human fMRI, the right primary visual cortex (V1) and right inferior temporal gyrus also showed greater activation to 2D versions of objects versus 3D objects (Snow et al. 2011). Our results also suggest that dorsal regions of the dog brain may process abstract features of object stimuli that include, but are not limited to, actions (Freud et al. 2017).

There were several limitations to our studies, the foremost being that only a subset of dogs participated in both the localizer study and the 2D-3D study. Some dogs were unavailable for both studies, and some dogs were unable to remain still while viewing video stimuli in the MRI. Further, we limited the number of objects to two or three items, which allowed for a simple controlled design with many trials per item but may limit the generalizability of our findings to all objects that a dog may encounter. Unlike human imaging studies, we did not include more abstract stimuli that were composed only of lines or were limited in color to black and white. To address these concerns, future research could confirm the selectivity of object processing regions for each dog using novel stimuli.

### Conclusions

Our fMRI results provide evidence for dedicated object processing regions in the occipital and parietal cortices in dogs. Although real objects and pictures of the same objects share a degree of visual similarity, they differ fundamentally in the actions associated with them and require experience wither either dimension. Further, even children at age 4 can show confusion about the properties of pictures and the objects they depict and the consequences of actions on pictures and objects (Ganea et al. 2009). We have begun to understand how dogs perceive their world through brain imaging, as this offers direct insight from the participant about the neural mechanisms underlying perception. Our studies reveal that there are potentially shared neural mechanisms underlying dogs’ and humans’ visual perception of objects, and that neural biases may in turn affect perception and behavior. These studies provide insight into the question of whether pictures are an appropriate proxy for real world stimuli for dogs and for fMRI.

## Ethics Statement

This study was performed in accordance with the recommendations in the Guide for the Care and Use of Laboratory Animals of the National Institutes of Health. The study was approved by the Emory University IACUC (Protocols DAR-2002879-091817BA and DAR-4000079-ENTPR-A), and all owners gave written consent for their dog’s participation in the study.

## Data Availability

The datasets generated during the current study are available from the corresponding author on reasonable request.

## Acknowledgments

Thank you to all of the owners who trained their dogs to participate in fMRI studies: Lorrie Backer, Rebecca Beasley, Emily Chapman, Darlene Coyne, Vicki D’Amico, Diana Delatour, Jessa Fagan, Marianne Ferraro, Anna & Cory Inman, Patricia King, Cecilia Kurland, Claire & Josh Mancebo, Patti Rudi, Cathy Siler, Lisa Tallant, Nicole & Sairina Merino Tsui, Ashwin Sakhardande, & Yusuf Uddin.

